# Novel Pan-ALDH Inhibitor KS100 Effectively Targets ALDH+/CD138⁻ Stem-like Cells to Overcome Relapse in Multiple Myeloma

**DOI:** 10.1101/2025.03.06.641909

**Authors:** Robert Chitren, Krishne Gowda, Shantu Amin, Gavin P Robertson, Subash Jonnalagadda, Tulin Budak-Alpdogan, Manoj K. Pandey

## Abstract

Multiple myeloma (MM), a clonal plasma cell disorder is the second most frequent hematological malignancy in the United States. This malignancy is characterized by a series of symptoms such as bone lesions, hypercalcemia, renal failure, and anemia. The current clinical drugs in the market are successful in treating multiple myeloma patients into remission but does not address relapse where a more aggressive phenotype of the cancer remains untreatable. We hypothesize that a small subset of multiple myeloma stem-like cells (MMSLC’s) that overexpress aldehyde dehydrogenases (ALDH^+^) is the cause of the relapse. Overexpression of ALDH bolsters drug resistance via detoxification and stemness via the retinoic acid signaling pathway. The phenotype of MMSLC’s is not yet known for certainty but there are a few well established markers such CD138 negative (CD138^neg^) cells that are known to overexpress ALDH. In this study, we target regular MM cells and bortezomib resistant ALDH^+^/CD138^neg^ MMSLC’s with a novel, potent, pan-ALDH inhibitor, KS100. Here we report KS100 effectively lowered ALDH expression in regular and bortezomib resistant ALDH^+^/CD138^neg^ cells, MM cell viability as well as proteins associated with MMSLC’s. Most importantly we showed that KS100 lowered ALDH^+^ populations in regular, bortezomib resistant and CD138^neg^ cells via ALDEFLUOR™ assay.

## Introduction

Multiple Myeloma (MM) is the second most frequent hematological malignancy of clonal plasma cells inside of the bone marrow [1]. These cells have many genetic abnormalities that affect prognosis and treatment. First-line treatment bortezomib [1], a proteasome inhibitor has shown initial success in effectively reducing cancer in patients where they will remiss, but acquired drug resistance makes MM incurable where all patients with acquire relapse and refractory disease [2].

Our hypothesis, and one that is most accepted of why relapse and refractory occur is that MM cancer stem-like cells (MMSLCs) exist in a small subset of the overall tumor population with properties of drug resistance, stemness and clonogenic regeneration [3]. These MMSLC’s survive initial conventional chemotherapeutic agents and then repopulate becoming the cause of tumor recurrence, therefore new drugs targeting MMSLC’s urgently need to be developed [4, 5].

The expression of aldehyde dehydrogenase (ALDH) is a critical factor in why MMSLC’s survive conventional chemotherapeutic agents and allow the cancer to repopulate [6]. The Aldehyde Dehydrogenases (ALDHs) are a superfamily that includes 11 families and a total of 19 members: ALDH1s (1A1, 1A2, 1A3, 1B1, 1L1, and 1L2), ALDH2, ALDH3s (3A1, 3A2, 3B1, and 3B2), ALDH4A1, ALDH5A1, ALDH6A1, ALDH7A1, ALDH8A1, ALDH9A1, ALDH16A1, and ALDH18A1 that all participate in critical cellular mechanisms such as aldehyde detoxification, lipid peroxidation, reactive oxygen species detoxification and retinoic acid synthesis [7, 8]. In a NAD(P)+-dependent reaction this family of enzymes oxidize endogenous and exogenous toxic aldehydes to less reactive, more biologically manageable carboxylic acids [9]. Also, ALDH playing a rate-limiting role in retinoic acid synthesis it promotes stemness by activating RAR/RXR transcription factors which transcribe proteins such as Notch, HOX, SOX2, SHH, Nestin, Akt, Pi3K, c-Myc, and RARβ [8, 10–14]. Consequently, cancer cells often overexpress specific ALDH isoforms to combat oxidative stress primarily due to high metabolic demands, radiation, and ROS-generating chemotherapeutics. Ultimately, overexpression of ALDH stem cell renewal, proliferation, differentiation, and anti-apoptotic properties ROS [11, 15, 16]

The identification of MMSLC’s has been a controversial hot topic on what should be the decided plasma B-cell phenotype. Some include CD138^neg^, CD19^+^/CD27^+^, CD38^high^/CD138^neg^ cells, and CD19^+^/CD20^+^/ CD138^neg^/CD38^neg^ cells [17, 18]. Our group agrees on using the CD138^neg^. The CD138^neg^ phenotype has been observed having elevated ALDH levels. ALDH-overexpressed (ALDH^+^) cells have greater tumorigenicity and chemotherapy resistance compared to ALDH^low^ cells [19]. We additionally use the CD138^neg^/ALDH^+^ phenotype due to the ability to measure cytosolic ALDH using an ALDEFLUOR™ assay [10, 20]. The inhibition of ALDH activity has growing evidence that it can be used as a potential strategy to eliminate/sensitize MMSLC’s preventing relapse and refractory from occurring [21]. Several ALDH inhibitors such as tetraethylthiuram disulfide (Disulfiram), 4-dimethylamino-4-methyl–pent-2-ynthioic acid-S-methylester (DIMATE), citral, and *N,N*-diethylaminobenzaldehyde (DEAB) have been developed but all either lack efficacy due to ALDH isoform specificity or are too toxic [8, 22, 23]. Unfortunately, there are currently no FDA inhibitors on the market. Thus, continued development of pan-ALDH inhibitors is needed.

To address these limitations, we explore the use of KS100, a novel, pan-ALDH inhibitor. This molecule contains an isatin moiety which earlier molecular docking studies have shown potently bind to ALDH1A, ALDH2 and ALDH3A1 active sites [6]. We chose ALDH1A, ALDH2 and ALDH3A1 in this as they span the three major ALDH families and are the most prevalent isoforms of those families to show the diversity and efficacy of KS100 as a pan-ALDH inhibitor. Previous studies have shown KS100 interacts with the W178 residue and an H-bond with the free amine group within the ALDH1A1 ligand– binding pocket. Similarly, KS100 had interactions with the F459 and F465 residues along with an H-bond interaction between the free amine group and L269 residue within the ALDH2 ligand–binding pocket. Further, KS100 interacts with the R292 residue and an H-bond interaction with the G187 residue in ALDH3A1 ligand-binding pocket. With the ability to bind to the pan-ALDH binding pocket we propose that the cells are eliminated through an accumulation of aldehydes that damages cells by forming protein adducts through non-enzymatic, covalent bonds with lysine, cysteine, and histidine residues, as well as through increased ROS generation and lipid peroxidation[6].

In this study we show KS100 as a pan-ALDH inhibitor that targets regular MM cells and ALDH^+^ MMSLC’s that are drug resistant to first-line MM treatment bortezomib. KS100 effectively reduces ALDH^+^ populations, pan-ALDH protein and gene expression, cell viability, stemness, clonogenic potential and tumor size while increasing endogenous ROS.

## Methods and Materials

### Western Blot Analysis

Whole cell extracts were created by subjecting the various cells to RIPA Lysis and Extraction Buffer (Thermo Fisher Scientific #89900) and 100x protease phosphatase inhibitor cocktail for an hour to lyse. The cell lysates were centrifuged at 15,000 rpm for 10 mins to remove insoluble material. Supernatant was collected and protein estimated. Lysate was prepped with dithiothretiol and NuPAGE dye (Thermo Fischer Scientific #NP0008) before resolved into 4-12% NuPAGE gel (Thermo Fischer Scientific #NP0321BOX). After electrophoresis, the proteins were electro-transferred to PVDF membranes (Thermo Fischer Scientific #88518), blotted with relevant antibody, and detected by enhanced chemiluminescence reagent and exposed to BIO-RAD ChemiDoc MP Imaging System. Using Gentle ReView™ Stripping Buffer (VWR, #19G0856497), several protein detections were done on the same membrane to conserve time and samples. We repeated all crucial blot experiments two to three times.

### Cell Lines and Reagents

Human MM cells MM.1R, MM. 1S, U266, RPMI 8226 cells were purchased from the American Type Culture Collection (ATCC, Manassas, VA). Bortezomib resistant cells MM.1S BTZ-R and U226 BTZ-R were a generous gift from Dr. Nathan Dolloff, The Medical University of South Carolina, Charleston, SC, USA. MM.1R, MM. 1S, and RPMI 8226 cells were cultured in RPMI 1640 growth media (Cellgro, Manassas, VA) supplemented with heat-inactivated 10% Fetal Bovine Serum (Sigma, St. Louis, MO), 100 I.U./ml penicillin, and 100 μg/ml streptomycin (Cellgro, Manassas, VA). U266 cells were cultured in RPMI 1640 with 15% FBS [24]. KS100 agent was synthesized and characterized at Penn State College of Medicine, Hershey, PA [6]. All other agents were procured from Selleckchem (Houston, TX).

### MTT Assay

To determine the potency of the drug, the cell lines were counted and split to a concentration of 5000 c/w in a round bottom 96-well plate and treated with various concentrations of the drug ranging from 0-25 µm. The MTT assay was ran for 72h with the dye being added with 3h left to go. The plates were centrifuged at 1000 rpm for 5 mins, then the supernatant was removed, and 100 µL of dimethyl sulfoxide (DMSO) was added to each well to dissolve the formazan crystals. The optical density was measured by delta value (560-630 nm) on a Tecan Infinite M1000 Plate Reader (Männedorf, canton of Zürich, Switzerland). The results were then analyzed and processed in GraphPad Prism.

### ALDEFLUOR™ analysis

To determine the effect of KS100 on CD138^neg^ cells, an ALDEFLUOR**™** (Stem Cell Technologies) assay was used. Cells were treated with various amounts of KS100 for 24h. Thereafter, cells were harvested and stained for ALDH following manufacturer’s protocol [25]. After staining, cells were analyzed using a S3e BIO-RAD cell sorter.

### Quantitative real-time polymerase chain reaction

Total RNA was extracted from 2×10^6^ cells using RNeasy plus kits (Qiagen, Valencia, CA) according to the manufacturer’s protocol. RNA was quantified using an ND-1000 spectrophotometer (Nanodrop Technologies, Wilmington, DE) and complementary DNA constructed as described [24]. Real-time polymerase chain reaction (PCR) experiments were conducted using an ABI-Prism 7700 Thermal Cycler and TaqMan Universal PCR Master Mix (Applied Biosystems, Foster City, CA). Specific primers were used for quantifying gene expression. The housekeeping gene glyceraldehyde 3-phosphate dehydrogenase (GAPDH) was used for all ΔΔCt calculations. Relative expression was calculated using software ExpressionSuite (Thermo Fischer Scientific) as described previously described [26].

### Annexin V Assay

Human MM cells were plated, treated, and incubated for 24h, and then tested for Annexin V Live Dead assay (Luminex Corporation, #MCH100105). Cells were stained with 1:1 ratio of Annexin V dye at room temperature for 20 mins in the dark following manufacturer’s protocol. Data was analyzed using Muse^®^ Cell Analyzer.

### Reactive Oxygen Species Assay

Cells were grown and treated with KS100 in a 3×10^4^ concentration per assay. The positive control used was a 250× ROS inducer chemical that was provided in the Reactive Oxygen Species (ROS) Detection Assay Kit (Abcam). The negative control was an unlabeled cell sample. The experimental controls were treated with a 1000× ROS label in the proper amount of ROS assay buffer that was provided. Manufacturer protocol was provided with kit and followed. Cells were imaged using a Leica fluorescent microscope and measured at Ex/Em 495/529 to determine change in fluorescence. Cells were also seeded at 2.5×10^4^ in a 96-well plate to obtain 70-80% confluency. Cells were then treated with ROS assay buffer, ROS label and incubated for 45 mins at 37^◦^C. Fluorescence was measured at Ex/Em 495/529 nm in the presence of compounds and controls as previously stated using a Tecan Infinite M1000 Plate Reader. Background was subtracted from results and processed using GraphPad Prism.

### Lentiviral Transfection ALDH1A1-Knockdown

The ALDH1A1 human shRNA lentiviral particle (OriGene #14846V) was used to transfect U266 cells to knockdown the expression of ALDH1A1. U266 cells were cultured to 40% confluence in 12-well plates, then 50 uL of virus solution (1×10^7^TU/mL) were mixed with 1 mL RPMI 1640 containing 15% FBS. Transfection efficiency was measured by GFP fluorescence microscopy at 72 hours post-transfection. Optimal concentration of puromycin was applied to screen transduced cells to obtain stable cell lines. Western blotting was used to determine ALDH1A1 expression levels in U266. Annexin V, as previously described, was used to determine KS100 efficacy on the ALDH1A1 cells.

### CD138^neg^ Cell Separation

Cells were treated with MACSprep™ Multiple Myeloma CD138 MicroBeads, human (Miltenyi Biotec, #130-111-744) following the manufacturers protocols. Once cells were treated with the magnetic microbeads, they were loaded into a MS Column (Miltenyi Biotec, #130-042-201) and were eluted through a MiniMACS™ Separator where the separate fractions were collected into media then used for down-stream use [24].

### Colony Forming Assay

Cells were dyed with trypan blue and manually counted using a hemocytometer and diluted to 1×10^4^ cells per well then treated with KS100 and in combination with other chemotherapeutic agents in varying concentrations with the appropriate media and left to incubated 24h at 37^◦^C. Control cells were not treated. Cells were then placed into a 1.5mL microfuge tube and centrifuged at 900 RPM for 5 mins. Supernatant was removed and re suspended in 10 μl of media to achieve final plating concentration of 1×10^4^ cells. MethoCult™ H4230 media (Stem Cell Technologies, #04230) supplemented with 15% FBS and 10% Gibco Phytohemagglutinin M form (PHA-M) (Thermo Fisher, #10576015) stimulated leukocyte conditioned medium was allowed to thaw to room temperature (15 - 25°C). Cells were then re-suspended and added to 1.1mL of Methocult H4330 media and vortexed. The cell mixture was allowed to rest for 5 mins to allow air bubbles to dissipate from the mixture. A sterile 3mL syringe fitted with a new 16 Gauge blunt-end needle was used to dispense the cell mixture into a 35mm petri dish. The medium was then distributed evenly across the dish. The culture dishes were then placed into a 245mm square dish with a 35mm dish with 3mL of sterile water. The dishes were then incubated at 37°C, in 5% CO2 with ≥ 95% humidity for 16 days. The dishes were then removed and imaged using a digital microscope. Images were then analyzed using manual counting of the colonies for a complete result.

### Toxicity and efficacy studies in mice xenograft model

Female NOD-SCID-IL2R gamma null (NSG) mice aged five weeks were procured from Jackson Laboratories and maintained and monitored in the animal research facility at Cooper Medical School of Rowan University (CMSRU), Camden, NJ. All xenograft investigations were conducted at CMSRU in accordance with the ethical rules provided by Rowan University’s own Institutional Animal Care and Use Committee (IACUC). Mice were subcutaneously injected into the right flank with 5 X10^6^ U266 MM cells in a 1:1 ratio of RPMI-1640 and Matrigel basement membrane matrix (Becton Dickinson, Bedford, MA) to create a human MM xenograft. When palpable tumors (volume ∼100 mm^3^) appeared about 10 days following injection, animals were randomly assigned into four groups of five mice each. To quantify tumor volume, digital caliper measurements of the longest perpendicular tumor diameter were taken as follows: 4/3 X (width/2)^2^ X (length/2). Every three days, the tumor volume and body weight were measured. The animals were euthanized at the end of the study, and the tumors were removed, weighed, and snap-frozen for future research.

### Statistical analysis

The statistical significance of the results was analyzed by unpaired two-tailed Student’s t-test or one-way analysis of variance (ANOVA), or two-way ANOVA using GraphPad prism software. Each graph represents the average of at least three replicates with error bars on the graph representing standard deviation* P ≤ 0.05, ** P ≤ 0.01, *** P ≤ 0.001, **** P ≤ 0.0001. ns: nonsignificant.

## Results

### KS00 inhibits the protein and gene expression of multiple aldehyde dehydrogenase isoforms in regular and bortezomib resistant MM cells

ALDH is an important enzyme that we hypothesize is leading to drug resistance in relapse and refractory patients. The levels of different ALDH isoforms were evaluated in five MM cell lines NCI-H929, MM.1S, MM.1R, RPMI8266 and U266. Western blot analysis of ALDH1A1, ALDH2, and ALDH3A1 in MM cell lines revealed that each cell line had different ALDH isoform expression levels (**Figure 1A**). Further examination showed that cell lines U266 and MM.1S had the highest total expression shown of the three ALDH isoforms (**Figure 1A**). Using this information, we treated U266 cells and MM.1S cells with increasing concentrations using KS100. KS100 effectively lowered the total expression of ALDH1A1, ALDH2, and ALDH3A1 in both cell lines in a dose dependent manner (**Figure 1B**). The same can be said for bortezomib resistant cells (BTZ-R) U266-BTZ-R and MM.1S-BTZ-R cells (**Figure 3B**). Though we did see reduced expression of ALDH2 in U266-BTZ-R we saw less in MM.1S-BTZ-R which may be explained by extreme overexpression of the enzyme (**Figure 3B**). To quantify the levels of ALDH enzymes on a gene expression level in the MM cell lines, quantitative real time-polymerase chain reaction was performed to evaluate the differential expression of all the three ALDH isoforms treated with concentrations of 0, 5, 10 μm of KS100 and normalized with GAPDH. We found that ALDH1A1, ALDH2 and ALDH3A1 were downregulated by KS100 in a dose dependent manner in all regular and BTZ-R cell lines (**Figure 1C,3C**). This is due to the feed-back loop mechanism that regulates ALDH. When ALDH is inhibited, retinal is not converted into retinoic acid. With retinoic acid not being produced, it then does not bind to the RAR/RXR co-activator complex (**Image X**) which activates transcription factors RARE/RXRE which transcribes ALDH [27].

**Figure 1:**
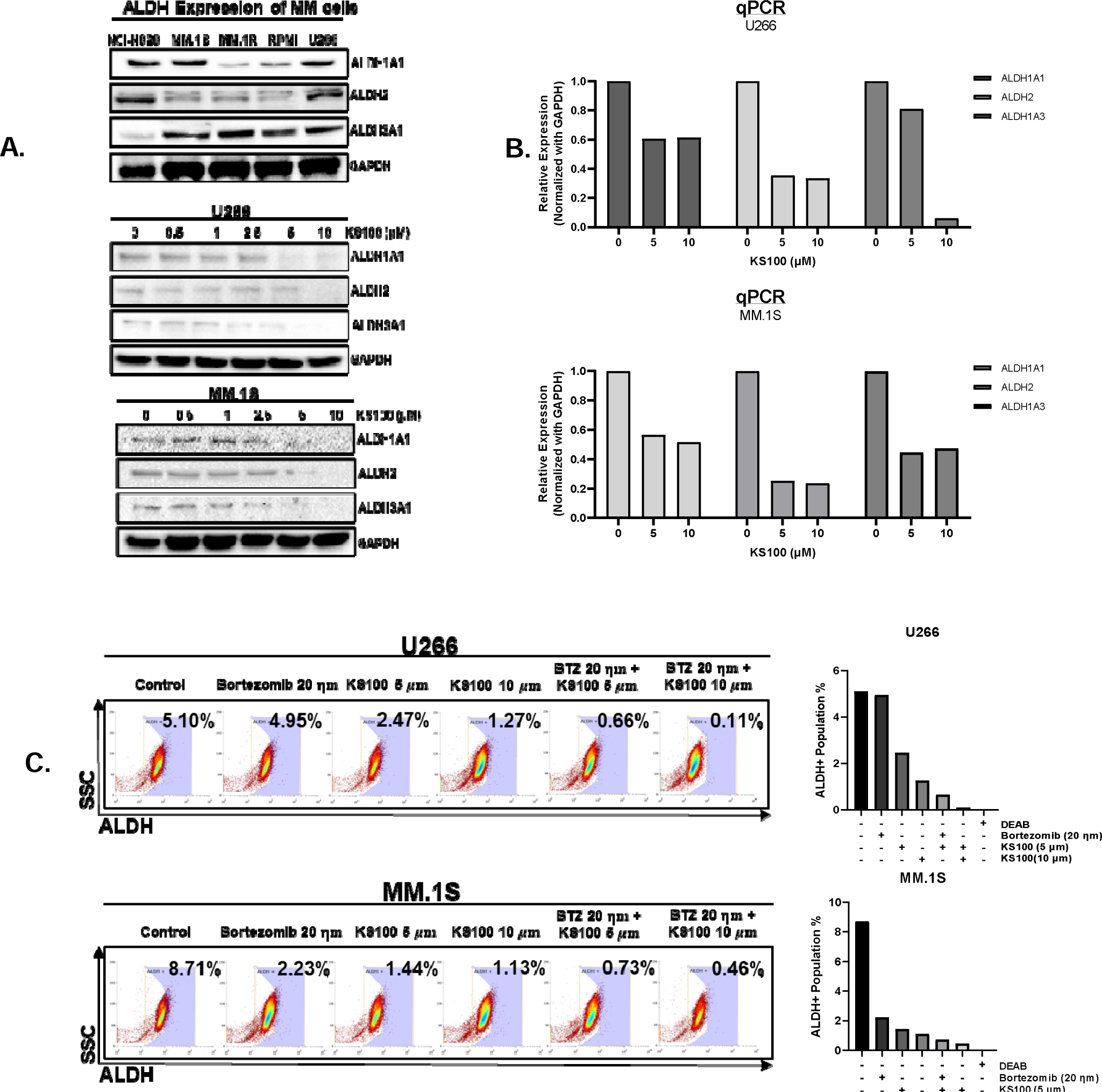
Effect of KS100 on ALDH expression and ALDH+ populations in regular MM cells. **A-B,** out of five MM cell lines U266 and MM.1S expressed the most ALDH1A1, ALDH2 and ALDH3A1. KS100 inhibits ALDH isoforms in a dose-dependent manner. U266 and MM.1S MM cells were treated with the indicated doses of KS100 for 24h. After incubation, the cells were lysed and immunoblotted with the mentioned antibodies. GAPDH antibody was used as a loading control after stripping and re-applying of the anti-bodies. **C**, Expression of BTK in various MM cell lines determined by real-time quantitative PCR (RT-qPCR). GAPDH was used as an internal control. **D-E** ALDEFLUOR™ assay demonstrating ALDH+ populations in U266 and MM.1S and the affect KS100 has on those populations.

### KS100 selectively targets ALDH in U266-ALDH1A1-knockdown cells

KS100 is a new novel, pan-ALDH inhibitor whose cytotoxic selectivity has not been explored yet. To test the selectivity of KS100, U266 cells was transfected with ALDH1A1 human shRNA lentiviral particles to knockdown ALDH1A1 expression. After this treatment, using GFP fluorescence microscopy we confirmed the cells were successfully transfected (**Figure 2A**). To confirm ALDH1A1 knockdown a western blot was performed compared to a GFP control. We found successful knockdown of ALDH1A1 expression in our transfected cells (**Figure 2B**). Further, we tested the selectivity of KS100 by treating our transfected cells with 5μm of the drug and performing an Annexin V apoptotic assay, comparing a U266-GFP-Control to our U266-ALDH1A1-Knockdown (**Figure 2C**). We found that compared to the control our U266-ALDH1A1-Knockdown cells when treated with KS100 mirrored the untreated cells leading us to say that KS100 is a highly selective ALDH inhibitor.

**Figure 2:**
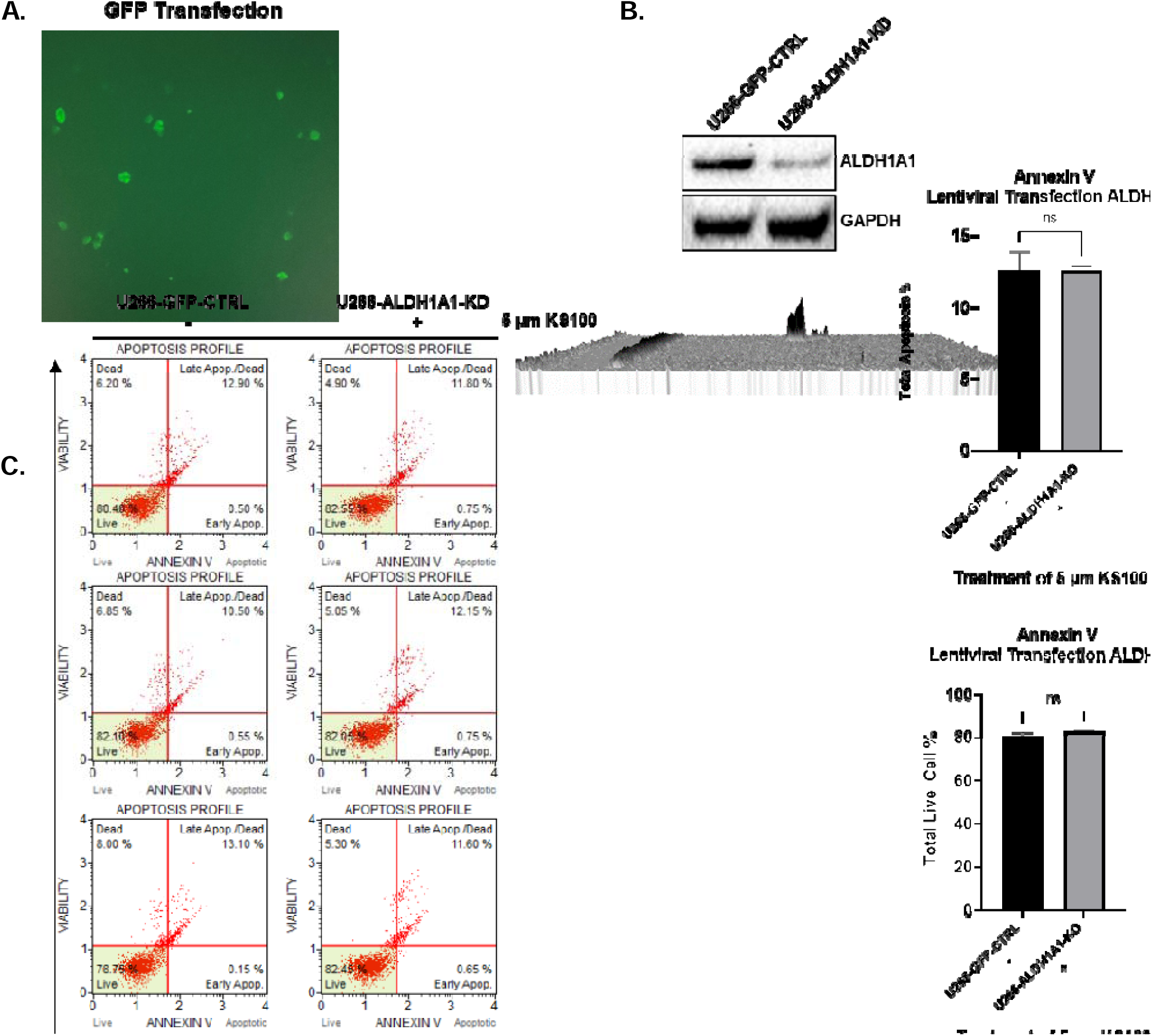
Selectivity of KS100 demonstrated by Lentiviral ALDH1A1-Knockdown Model. **A-B,** GFP and western blot confirmation of successful ALDH1A1-knockdown transfection in U266. Western blot U266-ALDH1A1-KD expression compared to U266-GFP-Control and GAPDH. **C**, Apoptotic cells are detected by Annexin V staining. U266-GFP-Control and U266-ALDH1A1 cells were treated for 24h with KS00 (5µM) alone then stained with Annexin V dye and analyzed using the Muse® Cell Analyzer. GraphPad prism was used for graphical representations and statistical analysis.

### KS100 reduces ALDH+ populations in regular, bortezomib-resistant and CD138^neg^ MM cells

The increased expression of ALDH in MMSLC’s give the cells an ability to detoxify themselves efficiently enough where current drugs on the market are ineffective in killing/reducing their numbers enough to prevent relapse and refractory in patients. Using an ALDEFLUOR™ assay we demonstrated the population of ALDH+ cells to be 5.10% of the total population in U266 and 8.71% of total population in MM.1S (**Figure 1D-E**). Both percentages fit in the range of what the previous research literature says [8, 28, 29] Treating U266 and MM.1S cells with an effective dose of 5μm of KS100 and then in combination with standard of care agent bortezomib, 20nm, demonstrated that our agent by itself and then in combination with bortezomib dramatically lowered the amount of ALDH^+^ MMSLC’s in the general population of cells. A previous study that characterized novel multi-isoform ALDH inhibitors identified KS100 to be the most potent pan-ALDH isoform inhibitor out of the series [16]. To showcase the drug’s ability to bind and inhibit ALDH expression an ALDEFLUOR™ assay was performed with regular U266 and MM.1S cells treated with current clinical agent bortezomib, KS100 and in combination. In Fig. untreated U266 cells express a population of ALDH+ cells of 5.10% which matches published literature values of 1-5% [18, 30]. Treatment of the currently clinically approved agent bortezomib had minimal effect on the population of ALDH^+^ cells we perceive to be MMSLC’s. As we start to treat with KS100 at 5 and 10 μm concentrations we see a 49.30% and a 24.90% drop in populations of ALDH^+^ MMSLC’s. When KS100 and bortezomib are used in combination nearly all ALDH^+^ cells are eliminated. The synergistic effect of these two drugs reduces the ALDH^+^ populations down to 0.66% and 0.11% of total cells. KS100 in combination with bortezomib at has dropped the population of ALDH^+^ MMSLC’s 88-98%. MM.1S expressed a population of ALDH^+^ cells at 8.71% of total population. A slightly different trend follows where we see a more drastic decrease of ALDH+ cells with the treatment of bortezomib followed by a lower decrease of ALDH+ at the 5um and10 um concentration which dropped the ALDH^+^ population to a mere 16.53% and 12.99% from its original population. When MM.1S is treated in combination the populations were reduced to 0.73 and 0.46% of total population (**Figure 1D-E**). When the same experiment was performed in U266 and MM.1S BTZ-R cells we found similar results. One different observation stood out for us in this experiment compared to the aforementioned. When BTZ-R cells are treated with bortezomib the ALDH^+^ populations increased compared to control cells. We hypothesize that when exposed to bortezomib these drug resistant cells have an acquired ability to increase the expression of ALDH to detoxify themselves from the drug (**Figure 3D-E**). This can be extrapolated into what we may see in a relapse and refractory patient and shows why pan-ALDH inhibitors such as KS100 are needed to reduce these overexpressed populations. Lastly, we explored the response of KS100 on BTZ-R ALDH^+^/CD138^neg^ populations. The definition of an MMSLC is a debated topic but most groups agree that CD138^neg^ cells have an overexpression of ALDH and are stem-like contributing to poor prognosis [31, 32]. We first separated cells using magnetic bead separation into CD138^pos/neg^ fractions and measured ALDH isoform expression using western blot (**Figure 5A**) and as expected CD138^neg^ fractions had higher expression of ALDH isoforms compared to CD138^pos^ cell populations. These MM BTZ-R CD138^pos/neg^ cell fractions were treated with KS100 and bortezomib. There are higher populations of ALDH^+^ cells in CD138^neg^ than CD138^pos^ in cells in both cell lines **Figure 5D)**. When treated with KS100 and with bortezomib we observed the same effect as MM BTZ-R cells (**Figure 3D-E**). When bortezomib is administered we observe the acquired ability of the cell lines to increase the amount of ALDH^+^ to detoxify themselves and then when administered KS100 the ALDH^+^ populations are dramatically reduced. Our hypothesis is that with KS100 we will be able to reduce the populations of ALDH^+^ MMSLC’s in drug resistant cancer to a level where relapse and refractory of MM will no longer be a threat.

**Figure 3:**
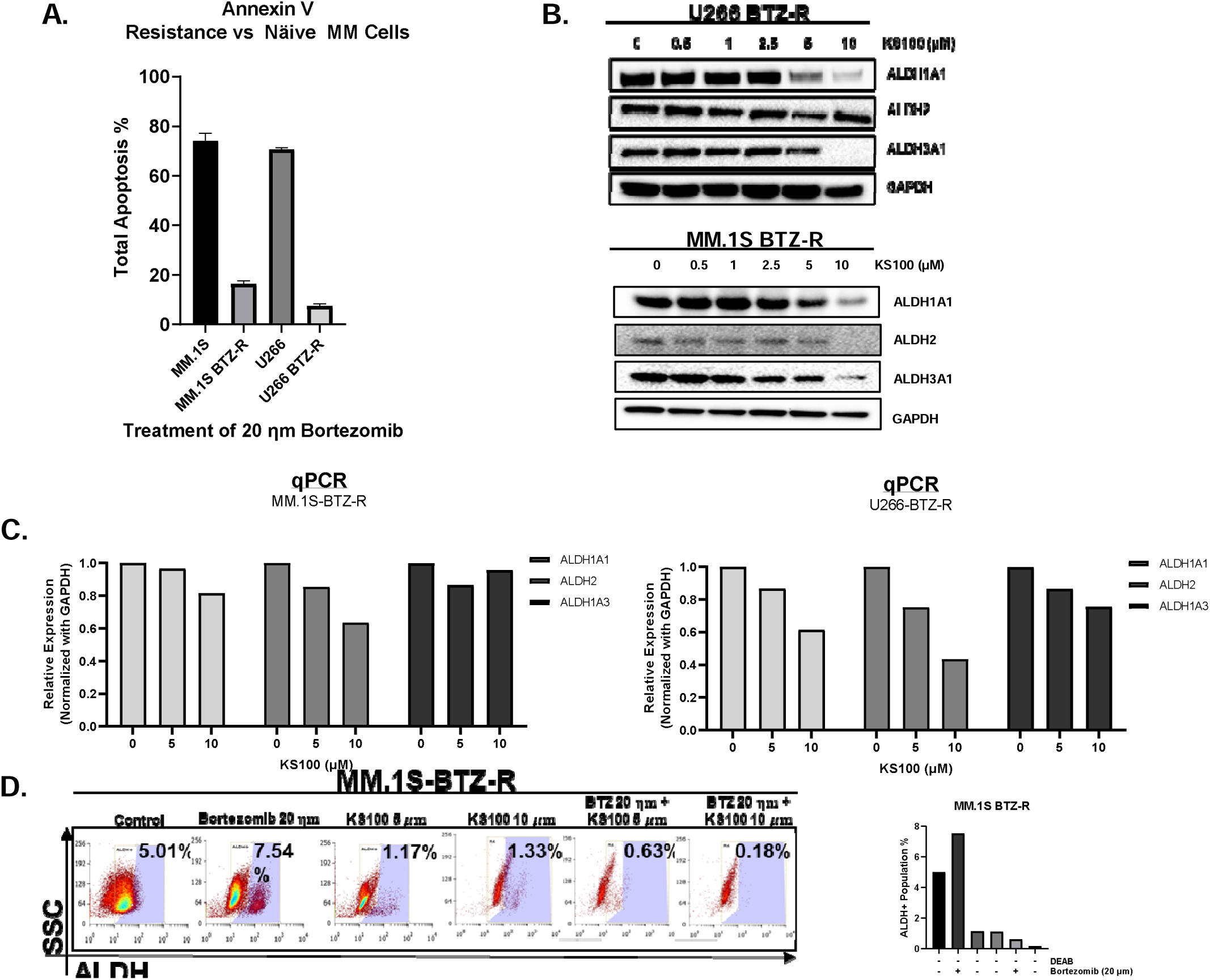
Effect of KS100 on ADLH expression and ALDH+ populations in bortezomib-resistant MM cells. **A,** Apoptosis measured via Annexin V of MM.1S and U266 regular cells compared to MM.1S-BTZ-R (bortezomib resistant) and U266-BTZ-R when treated with 20nM verifying drug resistant capabilities. KS100 suppresses ALDH isoforms ALDH1A1 and ALDH2 in U266-BTZ-R and all ALDH isoforms in MM.1S-BTZ-R in a dose-dependent manner. U266 MM cells were treated with the indicated doses of KS100 for 24h. After incubation, the cells were lysed and immunoblotted with the antibodies. GAPDH antibody was used as a loading control on the stripped membrane. **D-E,** ALDEFLUOR™ assay demonstrating ALDH+ populations in U266-BTZ-R and MM.1S-BTZ-R and the affect KS100 has on those populations.

### KS100 increases endogenous ROS generation within MM cells

ALDH enzymes are responsible for the oxidation of reactive aldehydes to their corresponding carboxylic acids. These aldehydes, if not detoxified, can react with cellular components, leading to oxidative stress and the generation of ROS. By inhibiting ALDH activity, the levels of reactive aldehydes may accumulate, promoting ROS formation, which can lead to cell death. We saw when MM-BTZ-R cells were treated with KS100 compared to our controls and untreated samples a 2-3× increase in reactive oxygen species (**Figure 4B)**. When cells experience a buildup of ROS, it can lead to oxidative stress, which is a condition characterized by an imbalance between the production of ROS and the ability of the cell to detoxify them or repair the resulting in damage or cell death.

**Figure 4:**
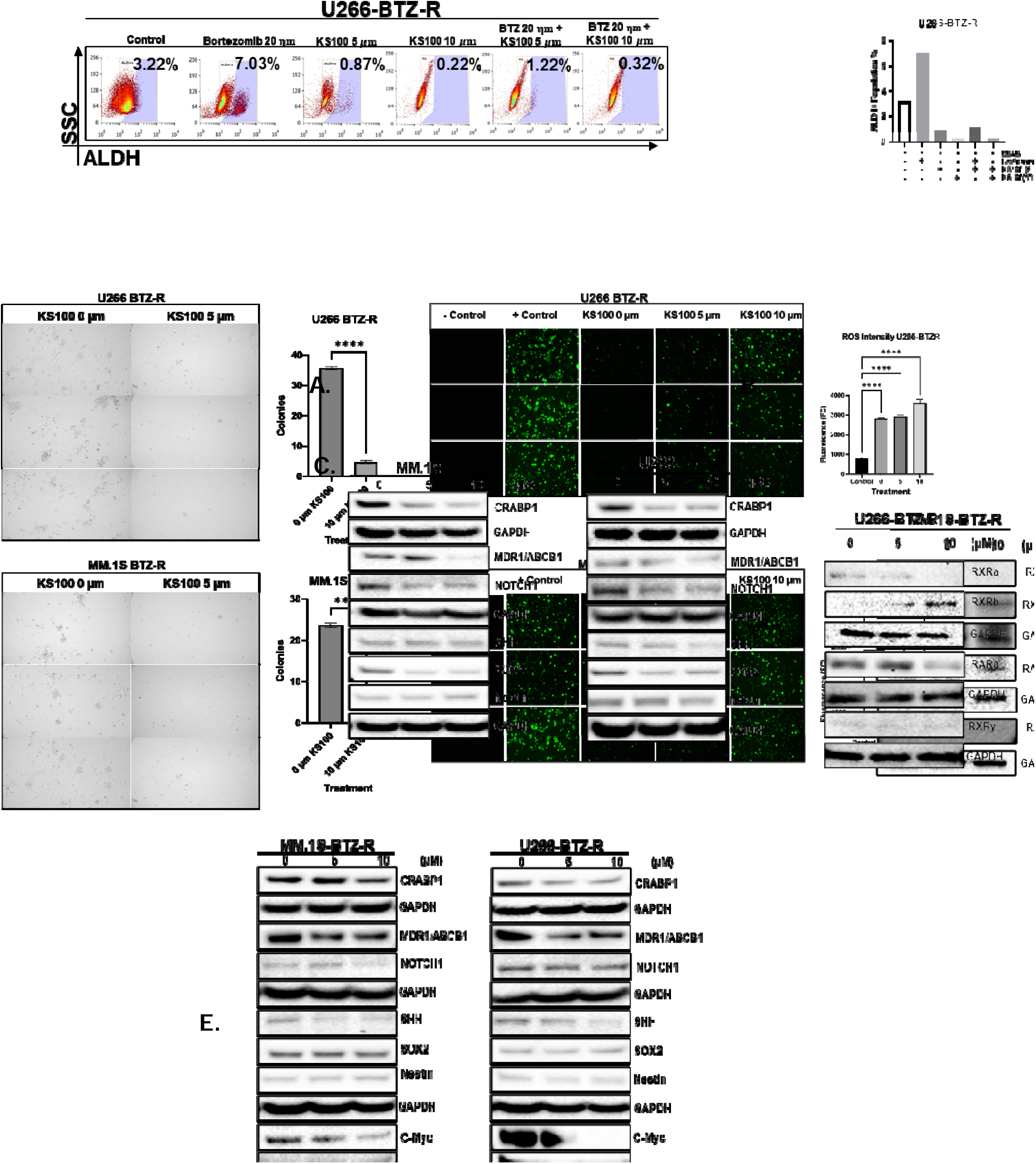
Effect of KS100 on clonogenic potential, ROS generation and stemness. U266-BTZ-R and MM.1S-BTZ-R cells were grown in colony supporting matrix for 14 days, with and without KS100 at a concentration of 5µM. Following the visual counting of colonies with a microscope, the survival fraction (SF) was calculated in relation to the untreated control group, and a graphical representation was made using GraphPad prism. **B,** Representative images of ROS fluorescence staining in U266-BTZ-R and MM.1S-BTZ-R cells treated with different concentrations of KS100 (5,10 µM). GraphPad prism was used to make graphical representations of the quantitative analysis of ROS fluorescence staining in MM-BTZ-R cells (n = 3). **C-E,** Western blot analysis of KS100 effect 92102on proteins associated with retinoic acid signaling pathway and stemness in U266, MM.1S, U266-BTZ-R and MM.1S-BTZ-R cells.

### KS100 reduces stemness via down-stream signaling of retinoic acid pathway

Retinoic acid, a metabolite of vitamin A, can upregulate the expression of ALDH1A1, ALDH1A2, and ALDH1A3 through activation of retinoic acid receptors (RARs) and retinoid X receptors (RXRs) [33]. This positive feedback loop contributes to the synthesis of retinoic acid and it’s signaling cascade. ALDH activity helps regulate the levels of retinoic acid within cells and tissues. By converting retinaldehyde to retinoic acid, ALDH enzymes control the availability of active retinoic acid for signaling purposes. Proper regulation of retinoic acid levels is essential for the precise control of downstream gene expression and cellular responses mediated by retinoic acid receptors [14, 34]. This crucial role in regulating stemness by influencing the expression and activity of various stemness-related proteins. Proteins such as HOX and Notch for differentiation, SOX2, SHH and Nestin for stem cell renewal, Akt and PI3K for apoptotic resistance and c-Myc for proliferation [35, 36]. Inhibition of ALDH by KS100 disrupts retinoic acid which in turn modulates downstream expression of various stem cell supporting proteins in regular and BTZ-R MM cells (**Figure 4C-E**). KS100 on top of inhibiting ALDH which leads to cell death or chemo-sensitization of bortezomib can also have negative effects on stemness by reducing stem cell supporting protein markers.

### KS100 reduces clonogenic potential in bortezomib resistant and CD138^neg^ MM Cells

The colony forming assay was employed to assess the survival and proliferative capacity of cells following treatment with KS100 compared to untreated control cells. Quantitative analysis of colony formation revealed a significant decrease in the number of colonies formed by treated cells compared to control cells. The mean number of colonies in the U266 KS100 treated group was 35 colonies, in the control group was 5 colonies. The mean number of colonies in the MM.1S KS100 treated group was 23 colonies, in the control group was 2 colonies (**Figure 4A**).Statistical analysis using unpaired T-test confirmed the significance of this difference (p < 0.0001). Representative images captured under a light microscope further illustrate the differences in colony formation between treated and control groups. Images of treated cell colonies depict sparse distribution and reduced colony density, whereas images of control cell colonies show dense packing and uniform distribution across the culture plate. KS100 effectively reduces the clonogenic potential in drug resistant ALDH^+^(**Figure 4A**). Furthermore, the same experiment was performed in CD138^pos/neg^ cell fractions as we found that bortezomib had very little effect on the CD138^neg^ cell population that we have proven contain overexpressed levels of ALDH. When CD138^neg^ cells are treated with KS100 by itself and in combination with bortezomib almost no colonies are observed compared to untreated control (**Figure 5C**). This observation leads us to say that KS100 effectively reduces clonogenic potential of drug resistant MMSLC’s which in turn can lower the probability of relapse/refractory.

**Figure 5:**
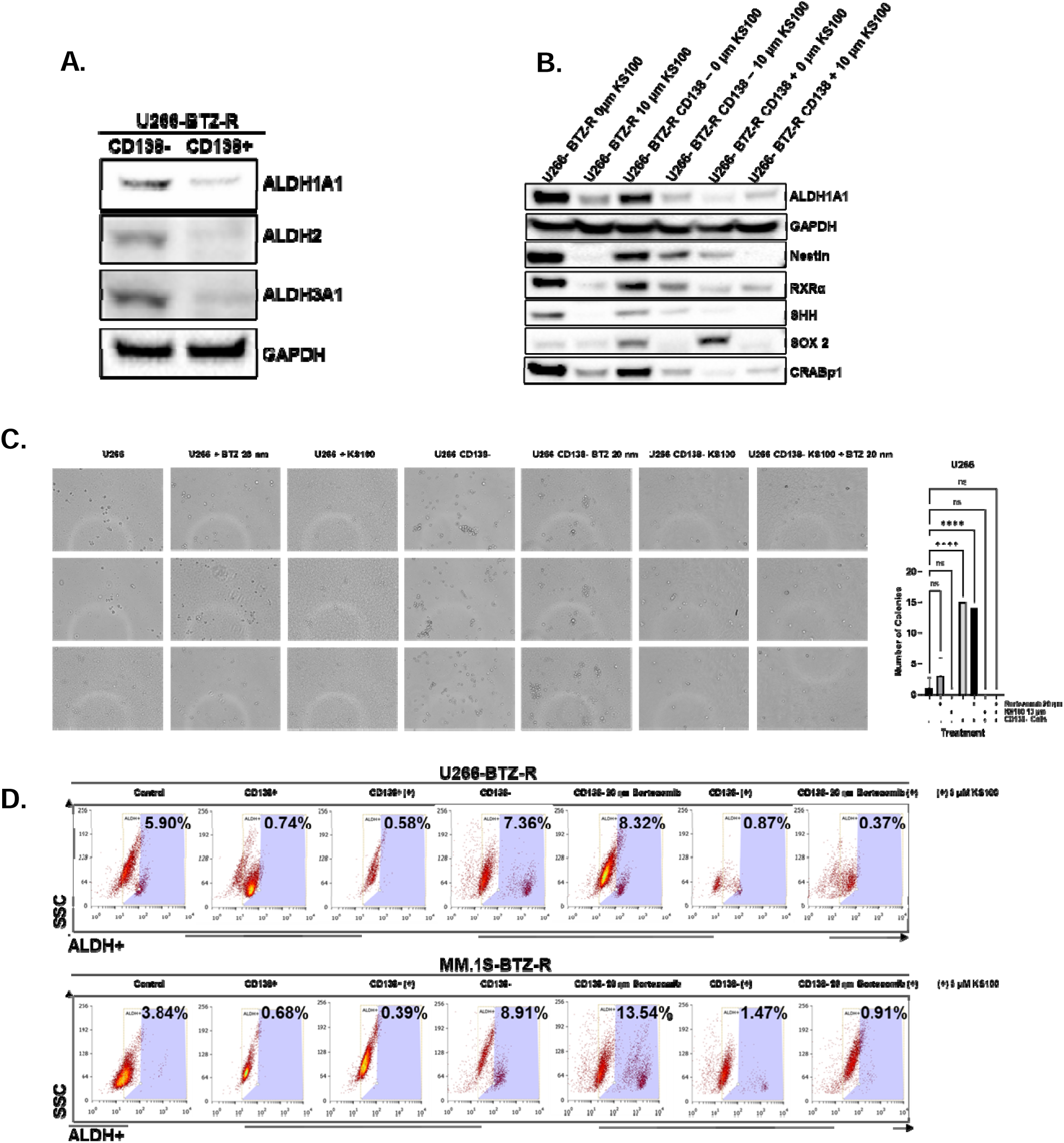
ALDH isoform expression and KS100 effects on clonogenic potential, stemness and ALDH+ populations in bortezomib resistant CD138-MMSLC’s. **A,** Expression of ALDH isoforms in CD138Neg cells fractioned from U266-BTZ-R cells via magnetic bead sorting. **B,** Western blot comparison of KS100 effect on proteins associated with retinoic acid signaling pathway and stemness in CD138-/+ populations fractioned from U266-BTZ-R cells via magnetic bead sorting. **C,** CD138-/+ populations were fractioned from U266-BTZ-R cells via magnetic bead sorting and were grown in colony supporting matrix for 14 days, with and without KS100 at a concentration of 5µM or bortezomib at 20nM. Following the visual counting of colonies with a microscope, the survival fraction (SF) was calculated in relation to the untreated control group, and a graphical representation was made using GraphPad prism. **D,** ALDEFLUOR™ assay demonstrating ALDH+ populations in CD138-/+ cells fractioned from U266-BTZ-R and MM.1S-BTZ-R via magnetic bead sorting and the affect KS100 and bortezomib have on those populations.

### KS100 shows promise in a model of MM xenografts as an pan-ALDH inhibitor

The MM xenograft model was developed by subcutaneous (S.C.) injection of U266 cells into the right flank of the NSG mice. The animals were randomized after approximately 10 days of (**Figure 6A**). We tested KS100s efficacy and safety in pilot tests and found that one weekly dose (q.wk) of 5 and 10 mg/kg was well tolerated. Two mice administered with 10 mg/kg was sluggish and hunched back on day 23, was euthanized. Importantly, all mice maintained normal weight compared to vehicle-treated mice (**Figure 6C**). Similarly, we examined KS100’s effectiveness in MM xenografted mice. The NSG mice were treated with 5 and 10 mg/kg, q.wk. We found that two doses of KS100 considerably reduced tumor burden in mice (**Figure 6B**). These data strongly imply we created a safe and highly effective ALDH inhibitor.

**Figure 6:**
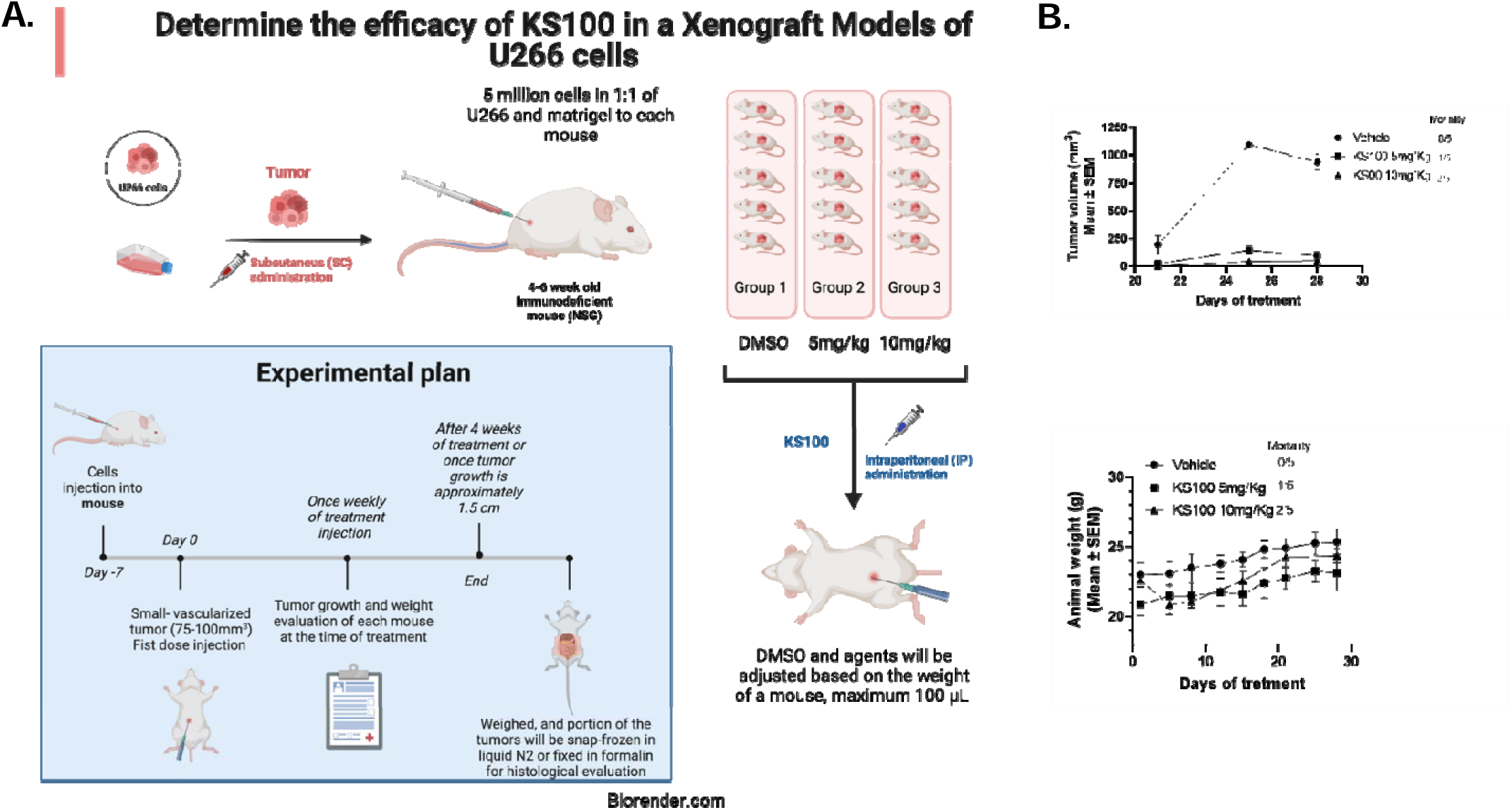
KS100 is safe and inhibits MM tumor growth. **A,** in a 1:1 mixture of RPMI 1640 medium and matrigel, five million cells were injected subcutaneously (SC) in the right flank of mice. Once tumor size reached approximately 100mm^3^, mice were randomly separated into three groups and treated with either DMSO or KS100 (5 and 10 mg/kg) intraperitoneally (IP) once a week (q.wk) for four weeks. **B,** KS100 is nontoxic to mice. Two KS100 doses were administered over the course of 28 days, and animal weights in xenografted mice were measured. On day 23, one sluggish mouse was euthanized. Presentation of animal body weight as mean ± SEM for vehicle (DMSO) and KS100 treated mice. **C,** Graph depicts tumor volume in mice treated with vehicle or KS00.

## Discussion

Multiple myeloma (MM) presents a complex therapeutic challenge, characterized by a high likelihood of relapse and drug resistance [37]. Despite initial responses to standard treatments like bortezomib, the emergence of refractory disease remains a significant clinical concern [38]. Understanding the mechanisms underlying MM relapse and identifying novel therapeutic targets are crucial for improving patient outcomes. Our study investigates the role of aldehyde dehydrogenase (ALDH) in MM relapse and refractory and its potential as a therapeutic target. ALDH enzymes play a critical role in detoxifying reactive aldehydes, thereby promoting cell survival and drug resistance in cancer cells, particularly cancer stem-like cells [15] .We hypothesized that targeting ALDH^+^ MMSLCs could effectively eliminate the population responsible for MM relapse. Our findings support this hypothesis, demonstrating that KS100, a potent multi-isoform ALDH inhibitor, effectively reduces ALDH expression in MM cells. This reduction in ALDH expression correlates with decreased cell viability via ROS accumulation, stemness markers, and clonogenic potential. Importantly, KS100 treatment leads to a significant reduction in ALDH^+^ populations, suggesting its ability to target MMSLCs in regular and drug-resistant cells. Moreover, our study elucidates the mechanisms underlying KS100’s anti-MM effects. We observed that KS100 enhances reactive oxygen species (ROS) generation within MM cells, leading to increased oxidative stress and impaired cell survival. Additionally, KS100 disrupts downstream signaling of the retinoic acid pathway, further attenuating stemness and tumorigenicity in MM cells. In a MM xenograft model, KS100 demonstrates robust efficacy in reducing tumor burden without significant toxicity. These preclinical findings underscore the potential of KS100 as a promising therapeutic agent for a relapse and refractory MM treatment. By targeting ALDH^+^ MMSLC’s, KS100 offers a novel approach to overcome drug resistance and prevent disease relapse in MM patients. Moving forward, clinical studies are warranted to further evaluate the efficacy and safety of KS100 in MM patients. Additionally, combination therapies incorporating KS100 with standard of care agents, such as bortezomib, merit further investigation to optimize treatment outcomes in relapse and refractory MM. Overall, our study highlights the therapeutic promise of targeting ALDH in MM and underscores the importance of developing innovative strategies to combat drug resistance and improve patient survival in this challenging disease.

## Author contributions

R.C.: Writing–review and editing, Writing–original draft, Validation, Methodology, Investigation, Formal Analysis. KG: Writing–review and editing, Validation, Methodology. SA: Writing–review and editing, Resources, Funding acquisition. SJ: Writing–review and editing, Resources, Funding acquisition. GR: Writing–review and editing, Resources, Funding acquisition. TB-A: Writing–review and editing, Resources, Funding acquisition. MP: Writing–review and editing, Writing–original draft, Visualization, Validation, Supervision, Resources, Project administration, Investigation, Funding acquisition, Formal Analysis, Conceptualization.

## FIGURE LENGENDS

**Supplementary 1:**
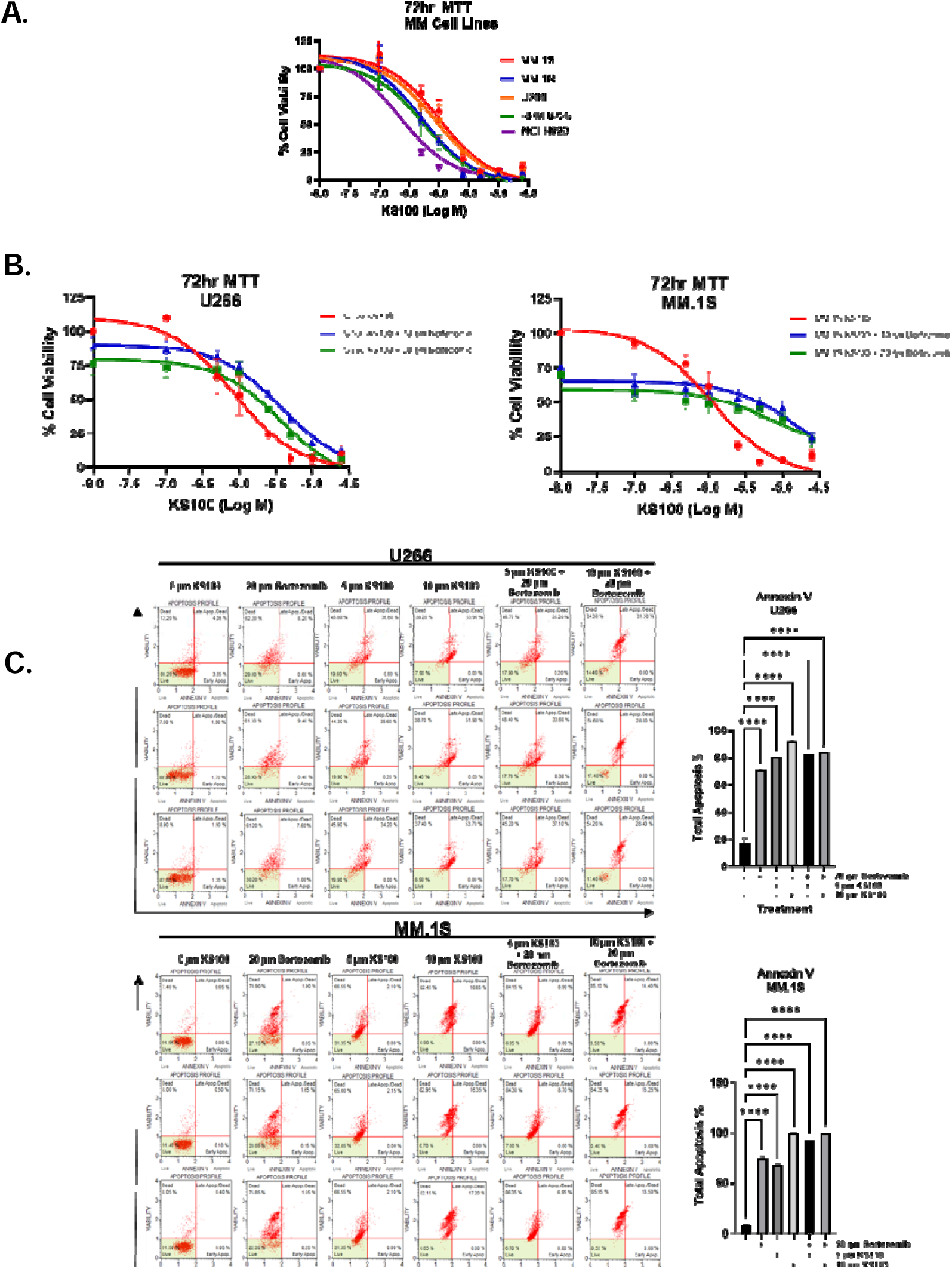
Cell Viability Data in MM Cells. The cytotoxic effects of KS100 were assessed using the MTT assay and compared to various chemotherapy agents. KS100 gradually reduced the viability of different multiple myeloma (MM) cell types, as illustrated in S1. A. After a 72-hour incubation, KS100-treated cells exhibited lower viability than those treated with other MM chemotherapeutic drugs in cell lines expressing ALDH (S1. B). Additionally, we conducted apoptosis assays on MM samples. Flow cytometry using Annexin V/PI staining revealed that KS100 treatment (5µM and 10 µM) for 24 hours significantly induced apoptosis in MM patient samples, resulting in an apoptosis rate of 92% (S1. C,D). We also examined the potential synergy between KS100 and 20 nM of bortezomib, a standard first-line treatment for MM. As shown in C,D, the combination therapy of KS100 with bortezomib led to a slight reduction in therapeutic efficacy. Specifically, the combined treatment of bortezomib (5nM) and KS100 (5µM) reduced MM cell viability by ∼84% (S1. C,D), suggesting that KS100 is itself more effective than bortezomib in reducing cell viability.

**Supplementary 2:**
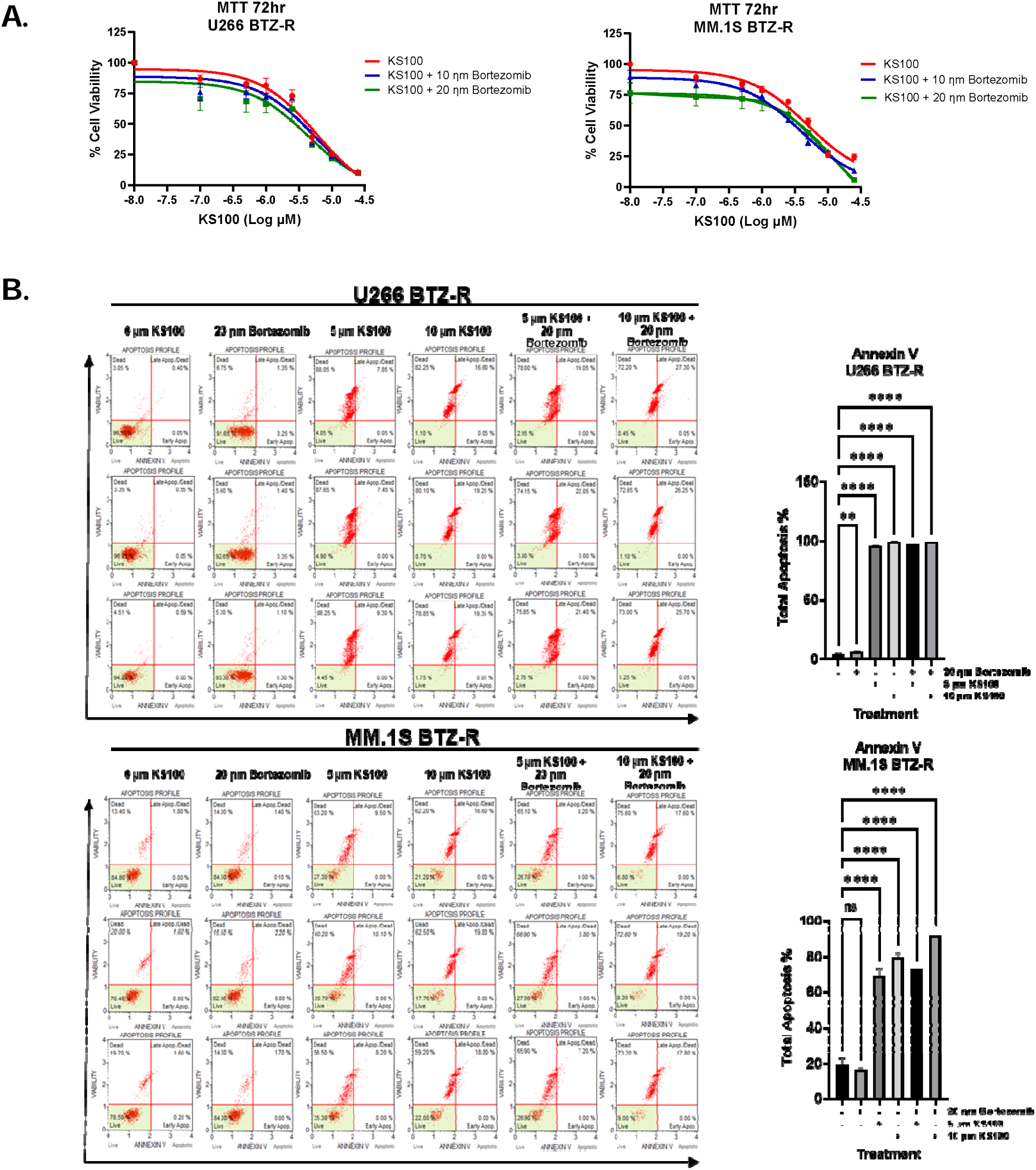
Cell Viability Data in Bortezomib Resistant MM Cells. The cytotoxic effects of KS100 were assessed using the MTT assay and compared to various chemotherapy agents. KS100 gradually reduced the viability of different multiple myeloma (MM) bortezomib resistant cell types, as illustrated in S2. A. After a 72-hour incubation, KS100-treated bortezomib resistant cells exhibited low viability than the resistant cells treated with bortezomib (S2. B). Additionally, we conducted apoptosis assays on MM samples. Flow cytometry using Annexin V/PI staining revealed that KS100 treatment (5µM and 10 µM) for 24 hours significantly induced apoptosis in MM patient samples, resulting in an apoptosis rate of 90-95% (S2. C,D). We also examined the potential synergy between KS100 and 20 nM of bortezomib, a standard first-line treatment for MM. As shown in C,D, the combination therapy of KS100 with bortezomib led to a similar response. Specifically, the combined treatment of bortezomib (20nM) and KS100 (5µM) reduced MM cell viability by ∼95% (S2. C,D), suggesting that KS100 is itself more effective than bortezomib in reducing cell viability.

